# Sharing happy stories increases interpersonal closeness mediated by enhancing interpersonal brain synchronization

**DOI:** 10.1101/2021.02.18.431771

**Authors:** Enhui Xie, Qing Yin, Keshuang Li, Samuel A. Nastase, Ruqian Zhang, Ning Wang, Xianchun Li

**Author notes:** These authors contributed equally (Enhui Xie, Qing Yin). Corresponding author: Xianchun Li, Shanghai Key Laboratory of Mental Health and Psychological Crisis Intervention, Affiliated Mental Health Center (ECNU), School of Psychology and Cognitive Science, East China, Normal University, Shanghai, 200062, China.

## Abstract

Our lives revolve around sharing stories with others. Expressing emotion (i.e., happy and sad) is an essential characteristic of sharing stories and could enhance the similarity of story comprehension across speaker–listener pairs. The Emotions as Social Information Model (EASI) suggests that emotional communication may influence interpersonal closeness, but the effect of sharing emotional (happy/sad) stories on interpersonal closeness remains poorly understood. Here, one speaker watched emotional videos and communicated the content of the videos to thirty-two listeners (happy/sad/neutral group). Both speaker and listeners’ neural activities were recorded using EEG. After listening, we assessed the interpersonal closeness between the speaker and listeners. Compared to the sad group, sharing happy stories showed a better recall quality and a higher rating of interpersonal closeness.

Meanwhile, the happy group showed higher interpersonal brain synchronization (IBS) in the prefrontal cortex (PFC) and temporal cortex (specifically TPJ) than the sad group. Moreover, such IBS mediated the relationship between the quality of sharing stories and interpersonal closeness, and happy emotion moderated this mediation model. The magnitude of IBS differentiated high interpersonal closeness from low interpersonal closeness. Exploratory analysis using support vector regression showed that the IBS could also predict the ratings of interpersonal closeness in left-out subjects. These results suggest that IBS could serve as an indicator of whether sharing emotional stories facilitate interpersonal closeness. These findings improve our understanding of sharing emotional information among individuals that guide behaviors during interpersonal interactions.

## Introduction

Sharing specific stories with another person plays an important role in social interaction. Sharing stories is a way for people to organize and convey their thoughts (Willems, Nastase, & Milivojevic, 2020), a way to enhance people’s ability to predict themselves and each other (Pickering, & Garrod, 2013), and a social practice promoting the formation of a collective memory (Hirst, & Echterhoff, 2012). Maswood and Rajaram (2019) have demonstrated that sharing stories is accompanied by expressing emotional meaning. Sharing an emotional story is a social interaction during which emotional brain states are transmitted between speaker and listeners (Chen et al., 2017; Hasson et al., 2012; Zadbood, Chen, Leong, Norman, & Hasson, 2017). For example, Zadbood and colleagues (2017) demonstrated that the listener mentally reconstructs the episodes of a story when listening to a speaker’s recollection of an audiovisual movie, even if the listener did not watch the movie before. Nonetheless, relatively little is known about how emotional states are proactively transmitted from speaker to listener.

The theory of Emotions as Social Information Model (EASI) proposes that expressing emotional information is a social signal in interpersonal interaction (Van, 2009). The listener receives both conscious and unconscious social cues from the speaker’s emotional expressions and the listener can regulate their emotional state to increase synchrony of emotional states with the speaker (Hari, Henriksson, Malinen, & Parkkonen, 2015). This alignment may influence interpersonal closeness of speaker and listener during emotion-related interaction. Previous studies have provided evidence that the expression of emotional information can promote mutual understanding, strengthen interpersonal communication, and promote social connections (Dubois et al., 2014; Smirnov et al., 2019).

Although according to EASI, emotion plays a crucial role in sharing stories and may influence interpersonal closeness, the definite effects of sharing positive and negative stories on influencing interpersonal closeness remain unevaluated. In the literature, two competing views have been presented. The first view is that sharing positive stories facilitates interpersonal closeness (Isgett & Fredrickson, 2015). When sharing emotional stories, individuals seem to prefer positive events and suppress negative events (Gillath, Bunge, Shaver, Wendelken, & Mikulincer, 2005; Piotroski, Wong, & Zhang, 2015). Initial behavioral studies have shown that participants preferred to post good-looking selfies, meaningful films, and music, and positive experiences on social media, thereby creating a favorable public impression (Dorethy, Fiebert, & Warren, 2014; Gable, Reis, Impett, & Asher, 2004; Pounders, Kowalczyk Christine, & Stowers, 2016). Sharing positive information can promote the attainment of desirable outcomes, such as obtaining social gratification from interpersonal interactions and strengthening interpersonal bonds (Fredrickson, 2001; Isgett & Fredrickson, 2015; Tamir & Robinson, 2007) by efficiently shaping a positive image to others (Johnson & Ranzini, 2018; Ranzini & Hoek, 2017). This suggests that sharing positive stories may play a more important role in enhancing interpersonal closeness. In direct opposition to the first view, another stream of research has suggested that sharing negative stories is critical to building a good interpersonal relationships (Shoham, Moldovan, & Steinhart, 2016); that is, sharing negative stories can enhance positive impressions and build close relationships based on powerful negative biases (Baumeister, Bratslavsky, Finkenauer, & Vohs, 2001; Fessler, Pisor, & Navarrete, 2015; Shoham, Moldovan, & Steinhart, 2016; Vaish, Grossmann, & Woodward, 2008; Rozin & Royzman, 2001).

The goal of current study is to clarify these evidently contradictory findings by exploring the influence of sharing stories with differing emotional content on interpersonal closeness. Previous neuroimaging studies have indicated that sharing a story can synchronize and align brain activity between speaker and listener (Dikker, Silbert, Hasson, & Zevin, 2014). For example, during speaker-listener communication several studies have observed alignment of neural responses between the speaker and listener in a network of high-level cortical regions typically attributed to the default-mode network, including the temporoparietal junction (TPJ), posterior medial cortex (PMC), and medial prefrontal cortex (mPFC; Stephens et al., 2010; Silbert et al., 2014; Zadbood, Chen, Leong, Norman, & Hasson, 2017). The similarity of the neural responses in speaker-listener dyads has also been associated with a shared interpretation of the narrative (Nguyen, Vanderwal, & Hasson, 2019; Zadbood, Chen, Leong, Norman, & Hasson, 2017). Further, emotional communication has been shown to increase the synchronization of brain activity across speaker and listener (Smirnov et al., 2019). Previous studies have shown that increased synchronization of brain activation in speaker-listener dyads was associated with a similar understanding of the emotional content and the similarity of their subjective feelings of valence and arousal (Nummenmaa et al., 2012; Smirnov et al., 2019). In sum, interpersonal brain synchronization (IBS) can be a neuromarker of various interpersonal interactions, including sharing stories and emotion communication (Nummenmaa et al., 2012; Stephens, Silbert, & Hasson, 2010). However, IBS as a neural indicator to uncover the neural mechanism of sharing emotional stories and interpersonal closeness during interpersonal interaction is yet unexplored.

The present study aims to provide behavioral and neural evidence for evaluating the effect of sharing emotional stories on influencing interpersonal closeness within the theoretical framework of EASI. Building on previous studies (Niedenthal & Setterlund, 1994; Ribeiro, Santos, Albuquerque, & Oliveira-Silva, 2019), the present study manipulates the valence of emotional stories (Happy vs. Sad) to reveal the effects of sharing emotional stories on interpersonal closeness. The neural mechanism of sharing emotional stories on influencing interpersonal closeness is investigated from the perspective of brain-to-brain coupling. On the behavioral level, we expected that sharing emotional stories would influence interpersonal closeness. Specifically, we hypothesized that sharing happy stories will increase interpersonal closeness more effectively than sharing sad ones. On the neural level, we expect that sharing happy stories will yield higher IBS than sharing sad stories. Finally, we hypothesized that enhanced IBS will mediate the effect of sharing emotional stories on influencing interpersonal closeness.

## Materials and Methods

### Participants

A total of 32 participants (age: 21.3 ± 2.4 years, 16 males) were enrolled as listeners in the present study. Inclusion criteria were (1) native Chinese speaker, (2) right-handedness, (3) normal speaking and hearing, (4) no previous or current neurologic/mental disorder according to self-report. Specifically, all the listeners were randomly assigned to listen the happy stories from the one competent speaker as speaker-listener dyads (15 listeners in the happy group) or the sad stories from the same competent speaker as speaker-listener dyads (17 listeners in the sad group).

One competent speaker (female, 19 years of age) was selected based on speaking sessions from 21 speakers. During this session, the speaker was asked to watch the emotional videos and narrate each video. From the 21 speakers, 16 were dropped due to failure in the comprehension test, 4 were dropped due to head motion and eye-movement artifacts. The narrations were recorded and qualitatively assessed with the understanding of the stories and the accuracy of the emotion in the stories by three independent raters. The brain data for the selected speaker were manually inspected for quality, and the data from the other speakers are not further analyzed here.

All participants provided written informed consent. The study had full ethical approval by the University Committee on Human Research Protection (UCHRP) University Committee on Human Research Protection (HR 403-2019), East China Normal University.

### Stimuli

The stimuli consisted of a total of 3 videos (happy video, sad video, and neutral video). The present study used 3 audio–visual movies, excerpts from the episodes of happy video (Hail the Judge, ~5-min in length), sad video (Root and Branches, ~7-min in length) and neutral video (natural scenes, ~6-min in length). These videos were chosen to have similar levels of production quality. Thus, each listener received two story stimuli, one neutral and one happy or sad. The average spoken recall recording of the happy story was 2.4 min (min: 1.7, max: 4.4) in duration, comprising 227 words; the average spoken recall recording of sad story was 4.6 min (min: 1.6, max: 12.8) in duration, comprising 453 words; and the average spoken recall recording of the neutral story was 3.5 min (min: 1.3, max: 7.4) in duration, comprising 380 words. Audio recordings were obtained from each speaker who watched and recounted the 2 videos (one neutral and one happy/sad) with EEG-recording. The listener listened to 2 corresponding audio recordings (Fig. 1B).

**Figure 1.**
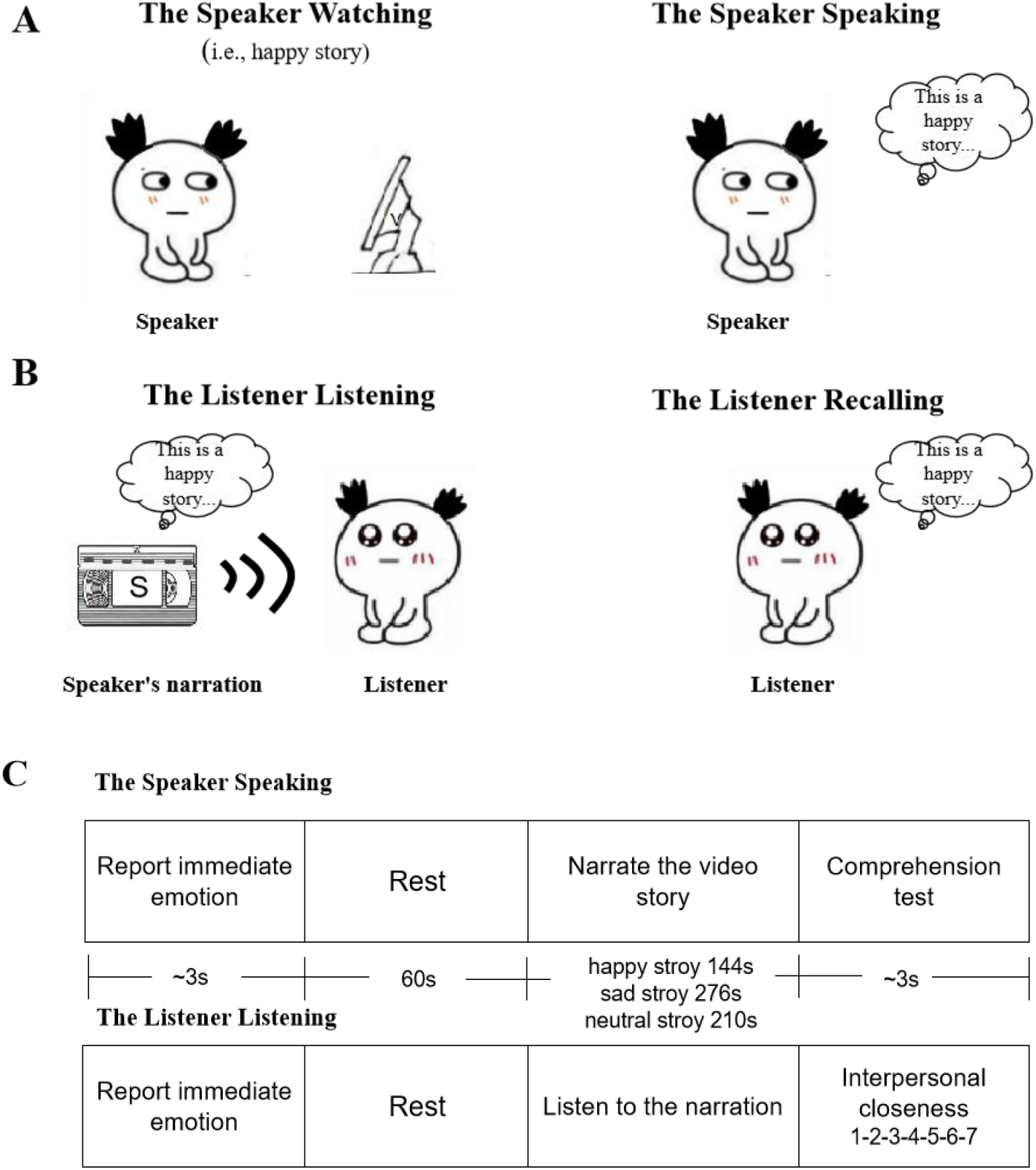
Experimental design. (A) Speaker design. The speaker was invited to watch an emotional video and shared the stories in the video by narrating. (B) Listener design. The listener was asked to listen to the story of the video through the speaker’s narration and allowed to recall the story which the speaker shared. (C) The task in The Speaker Speaking and The Listener Listening. The specific procedure of the sharing stories.

### Procedures

The experimental procedures consisted of a resting-state phase and a task phase for both the speaker and the listener sessions. The speaker and the listener performed their tasks separately. In all experimental stages, the neural activity of the speaker and the listener was recorded with EEG. During the resting-state phase, participants were instructed to relax while keeping their eyes closed without falling asleep and to avoid excessive head motion. For each dyad, an initial resting-state session served as baseline.

The task phase included two main sessions. In the first session (the speaker session), the speaker session included two main stages: in the first stage (Speaker Watching), the speaker participants were asked to watch the emotional (happy, sad, and neutral) videos (Fig. 1A); in the second stage (Speaker Speaking), the speakers were asked to verbally narrate the stories in the videos and their narrations were recorded (Fig. 1A). The speaker participants’ brain activity was recorded using EEG during speaking. Finally, one speaker was determined as a competent speaker. The recording from the competent speaker was listened to by all listeners, yielding 32 speaker-listener dyads. Finally, one competent speaker was chosen and fixed from recruited 21 speakers indicated by our comprehension test (Fig. 1C). A comprehension test was conducted to choose a competent speaker. Three independent raters listened to these audio recalls and assigned memory scores to each participant. Suggested items to consider were (a) understanding of the stories, (b) expression of the episodes, (c) the number of scenes remembered, (d) details provided, and (e) the accuracy of emotion in the stories. They reported a score for each participant and these numbers were rescaled (to have the same minimum and maximum) across the three raters.

In the second session (the listener session), a separate group of subjects were invited to listen to the emotional (happy/sad) and neutral recordings (The Listener Listening, see Fig. 1B) and recall the corresponding recordings (The Listener Recalling, see Fig. 1B). To control confounding effects of between-group differences in mood, all listeners were required to report their emotional state immediately before listening. The happy group received happy stories from the competent speaker’s recording, whereas the sad group received sad stories from the competent speaker’s recording. Moreover, sharing neutral stories served as a baseline for a sharing emotional performance and therefore it was reasoned that this condition should be performed before the happy or sad condition. To determine the effect of sharing emotional stories on interpersonal closeness, corresponding indices were assessed by means of the inclusion of other in the self (IOS) Scale (Aron, Aron, & Smollan, 1992) before recalling (Fig. 1C).

### EEG data acquisition

The neural activity of each participant was simultaneously recorded with an EEG recording system using a 64-channel system (Compumedics NeuroScan) with a sampling rate of 1000 Hz. The electrode cap was positioned following the standard 10-10 international system. Two vertical and two horizontal Electro-Oculogram (EOG) were placed to detect eye-movement artifacts. Impedances were maintained below 10 kΩ.

## Data analysis

### Behavioral data analysis

The quality of communication between speaker and listener was evaluated using The Listener Recalling stage, in which listeners were asked to recall everything they remembered from the stories they heard. Quality of recall was assessed by three raters (following the procedure in Zadbood et al., 2017). The raters first established the rating system by which the quality is principally judged by the detail level of the scene and the accuracy of the narration. Based on this system, they then rated all three stories from the listener independently on the same scale (from 0 to 30). The final quality score of each story was determined by averaging the three raters’ scores on this story. The primary behavioral index “delta recall quality” was computed in the following way: delta recall quality = average recall quality in the emotional group (happy or sad) – the corresponding neutral recall quality. That is, the score of the neutral memory served as a baseline, such that the final scores of emotional memories were subtracted by the mean score of the neutral memories. To evaluate the difference of the behavioral index in sharing quality between happy and sad groups, we conducted an independent-sample *t*-test. Cronbach’s alphas of 0.91 for the happy video, 0.92 for the sad video, and 0.94 for the neutral video indicate high consistency between the raters. An example of a rating sheet made and used by one of the raters is provided in Supplementary material (Table S1).

### EEG data analysis

The EEG raw data were preprocessed and analyzed using the EEGLAB toolbox (version 14.1.0; Delorme & Makeig, 2004) and in-house scripts in MATLAB (R2014a, The MathWorks, Inc.). EEG data were filtered with a bandpass ranging from 1 to 45 Hz and notch filter at 50 Hz. Data were re-referenced offline to an average of the left and right mastoid and downsampled to 250 Hz. EEG data were divided into consecutive epochs of 1,000 ms. Eye-movement artifacts were removed with an independent component analysis (ICA) method (Makeig, Bell, Jung, & Sejnowski, 1996). Signals containing EEG amplitudes greater than ± 75μV were excluded.

The subsequent data were submitted to an interpersonal brain synchronization (IBS) analysis known as phase locking value (PLV; Lachaux, Rodriguez, Martinerie, & Varela, 1999; Czeszumski et al., 2020). PLV was computed for each pair (i, k) of electrodes for each frequency band according to the following:

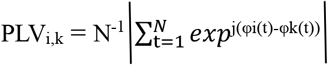

where N represents the number of trials, φ is the phase, | | represents the complex modulus, and i and k indicate the electrode from participants 1 and 2 in a dyad, respectively, where one participant is the speaker and the other is a listener. The PLV ranges from 0 to 1 where PLV equals 1 if the two signals are perfectly synchronized and equals 0 if the two signals are unsynchronized. Phases were extracted using the Hilbert wavelet transform (Schölkopf, Platt, Shawe-Taylor, Smola, & Williamson, 2001), and four frequency bands: theta (4–7 Hz), alpha (8–12 Hz), beta (13–30 Hz), and gamma (31–48 Hz) were identified as typical frequency ranges in previous studies (e.g., Delaherche, Dumas, Nadel, & Chetouani, 2015; Hu et al., 2018). Previous neuroimaging work has shown that brain activity in the theta band was correlated with emotion, memory encoding, and information transmitting (Ding, 2010; Klimesch, Doppelmayr, Russegger, & Pachinger, 1996; Symons, Wael, Michael, & Kotz, 2016; Zheng, Quan, & Zhang, 2012).

The present study calculated a delta PLV value in the theta band for each speaker-listener dyad using the equation “delta vPLV = average theta band PLV in the emotional group (happy or sad) – the corresponding neutral average theta band PLV”. We conducted independent-sample *t*-tests (Happy vs. Sad) on the IBS of speaker-listener dyads to explore the difference of the IBS in sharing stories between happy and sad groups. Differences were considered significant using an electrode-pairs-level threshold of *p* < 0.05 (Bonferroni-corrected).

We explore three possible elements of sharing emotional stories (Jeffres & Lin, 2006; Lasswell, 1948): the receiving matchup (The Speaker Watching-The Listener Listening), the sharing matchup (The Speaker Speaking-The Listener Listening), and the sending matchup (The Speaker Speaking-The Listener Recalling). In the receiving matchup, both speaker and listener are receiving information from either media or other people. During this matchup, the speaker received the content of emotional stories from the video stimulus and the listener received information about emotional stories from speaker’s narration. In the sharing matchup, the speaker is actively transmitting the content of the emotional stories to listeners via verbal and the listeners are listening to speaker’s narration. In the sending matchup, both the speaker and listener are transmitting information to others via verbal recall. The speaker is transmitting information to listeners and the listeners are verbally recalling the information they received from the listener. To further explore whether the delta value of the IBS was strongly associated with sharing emotional stories, several validation analyses were performed. Compared with the receiving and sending matchups, stronger IBS (delta value) were observed in the sharing matchup, so subsequent data analysis focused on the sharing matchup (The Speaker Speaking-The Listener Listening; Ahn et al., 2018; Chen et al., 2020). A subsequent correlation analysis examined the association between the behavioral (delta recall quality) and neural index (delta PLV in The Speaker Speaking-The Listener Listening).

### Moderated mediation model

Moderated mediation analysis was conducted using the SPSS macro PROCESS (Hayes, 2013) to reveal the underlying processes that may mediate or moderate the relationship between sharing emotional stories and interpersonal closeness. The quality of recall (delta value) served as the independent variable, the delta value of the IBS as the mediator, and the interpersonal closeness as the dependent variable in mediation model to reveal the relationship between sharing emotional stories and interpersonal closeness. Moreover, the valence of the emotion was hypothesized to play a moderating role in the association between the value quality of recall and the IBS.

### Coupling directionality

To estimate the information flow between the speaker and the listener during the Speaker Speaking-Listener Listening matchup, Granger causality (G-causality) analysis was conducted. According to Granger theory (Granger, 1969), for two given time series X and Y, if the variance of the prediction error for the time series Y at the current time is reduced by including historical information from the time series X in the vector autoregressive model, then the changes in X can be understood to “cause” the changes in Y. The MVGC Matlab toolbox (Barnett & Seth, 2014) was used to estimate the full multivariate conditional G-causality. The task-related data for each participant were z-scored prior to G-causality analysis based on the mean and standard deviation of the resting-state signal. Further, the paired-samples *t*-test was used to compare the difference between two directions of G-causality.

### Prediction of interpersonal closeness

We conducted a discriminative analysis to test whether the IBS of the speaker-listener dyads could discriminate interpersonal closeness. First, several lines of research have proposed that IBS can be used as classification features (Hou, Song, Hu, Pan, & Hu, 2020; Pan et al., 2020). The interpersonal closeness scores were dichotomized into high and low groups via the median split. We used all electrode pairs as the features to classify the high and low interpersonal closeness, rather than just significant pairs, to avoid training biases of the feature extraction. A support vector classifier (SVC) based on support vector machines (SVM) was used to classify high and low interpersonal closeness based on IBS of the speaker-listener dyads. The LIBSVM toolbox (http://www.csie.ntu.edu.tw/~cjlin/libsvm, Chih-Chung, Chang, Chih-Jen, & Lin, 2011) was used to conduct SVC analysis. For cross-validation, the classier was trained on the IBS from all electrode pairs in a randomly selected 70% of the data. Subsequently, we tested the model on the left-out 30% of all datasets. Thirdly, the training dataset was trained by nu-support vector classifier (V-SVC) with the radial basis function (RBF) and nu was set to the default of 0.5. The other two parameters (C, γ) were used to adjust the efficiency of the algorithm and the best parameters (C, γ) were learned via a 5-fold cross-validation hyperparameter grid search within the training set (Yan, Wang, & Cai, 2008). Finally, based on the probability estimates from the SVC model, the area under the receiver operating characteristic curve (AUC) was calculated (Faraggi & Reiser, 2002). Previous work has shown that AUC can effectively quantify the accuracy of the prediction based on synchronized brain activity (Cohen et al., 2018. The significance level (threshold at *p* < 0.05) was calculated by comparing the AUC from the correct labels with 2,000 randomization samples with happy/sad labels shuffled.

The classification analysis described above was used to explore if IBS can successfully be used to classify high and low interpersonal closeness in the present study. To support this discriminative analysis, we also conducted an exploratory support vector regression (SVR) analysis, a regression method based on SVM, to explore whether the IBS could predict interpersonal closeness (using the LIBSVM toolbox, Chih-Chung et al., 2011). Unlike the discriminative analysis, the response variable here was the listeners’ rating score of the interpersonal closeness. IBS from all electrode pairs was used as the features and a random 70% of the dataset was assigned as a training dataset and the remaining 30% of the data were used as a testing dataset in the same way as the prior analysis. The model was trained using ε-support vector regression (ε-SVR) with RBF. Based on Hou et al., (2020), the parameter ε was set to 0.01. The other two parameters (C,γ) were optimized using grid search via a 5-fold cross-validation method in the training set (Yan et al., 2008). Finally, the Pearson correlation coefficient indicated the prediction accuracy between the actual and predicted values (Kosinski, Stillwell, & Graepel, 2013). Collectively, these analyses were conducted to explore whether the IBS of the speaker-listener dyads could predict interpersonal closeness and further to test the generalizability and replicability of our results.

## Results

### Behavioral results

Examination of the recall quality from the happy and sad group revealed that significant better quality in the happy group compared to the sad group (*t* (30) = 4.14, *p* < . 001, Cohen’s *d* = 1.45, independent-sample *t*-test, see Fig. 2A).

**Figure 2.**
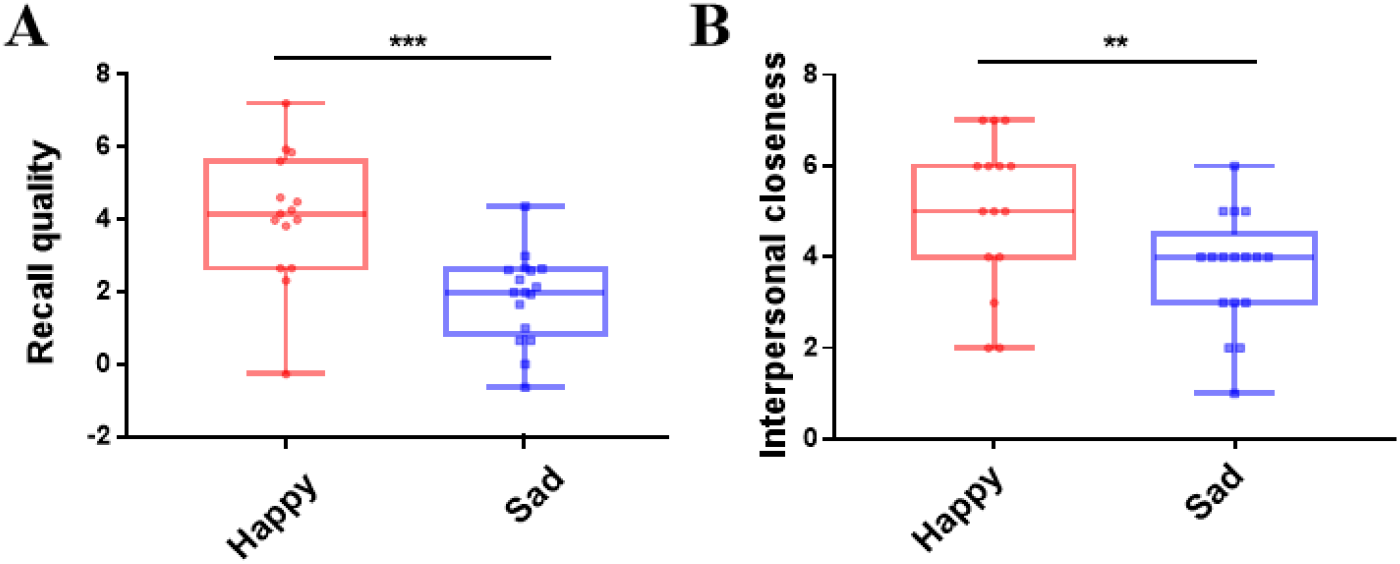
Behavioral results in the happy and sad group. (A) The recall quality (delta value) is shown for happy (Happy-Neutral) and sad (Sad-Neutral) emotions. (B) The rating of interpersonal closeness is shown for happy (Happy-Neutral) and sad (Sad-Neutral) groups. \*\**p* < 0.01, \*\*\**p* < 0.001.

In addition, we conducted an independent-sample t-test on the interpersonal closeness scores as measured by the IOS. Specifically, the result showed that the happy group had significantly higher scores than the sad group, *t* (30) = 2.91, *p* < 0.01, Cohen’s *d*= 0.94, see Fig. 2B.

### IBS of speaker-listener dyads

The present study compared the IBS in the theta band with other bands. The results showed more significant electrodes pairs were observed in the theta-band than other bands (all Bonferroni corrected, see details in the Supplement (Fig. S1A-S1D). Based on this result and previous research (e.g., Symons, Wael, Michael, & Kotz, 2016; Zheng, Quan, & Zhang, 2012), the theta band was used as the band of interest.

In the theta band (4–7 Hz), we found that the IBS was significantly higher in the happy group than that in the sad group (*t* (30) = 7.38, *p* < . 001, Cohen’s *d* = 2.69, Bonferroni corrected, the independent-sample *t*-test, Fig. S2). Further, 12 pairs had significantly higher PLV in the happy group than in the sad group (*t*s > 3.51, *p*s < . 001, Bonferroni corrected; detailed findings are shown in Table S2). The results indicate that the significantly increased IBS between dyads was specific to happy stories (Fig. 3). Moreover, PLV was highest in frontal and temporal regions for the speaker-listener dyads during shared emotional stories.

**Figure 3.**
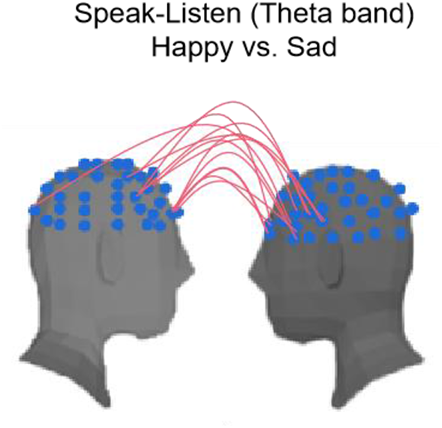
Significant electrodes pairs in theta band. PLV in theta band is shown that for happy (Happy-Neutral) and sad (Sad-Neutral) emotions, the significantly increased IBS between dyads was specific to happy stories in sharing matchup (The Speaker Speaking-The Listener Listening) after Bonferroni correction.

Further, the sharing matchup (The Speaker Speaking-The Listener Listening) was compared to the receiving matchup (The Speaker Watching-The Listener Listening) and sending matchup (The Speaker Speaking-The Listener Recalling) using two paired-sample *t*-tests (Fig. S2). Results indicated that the sharing matchup IBS was significantly higher than the receiving matchup IBS (*t* (31) =6.46, *p* < . 001, Cohen’s *d* = 1.19, Bonferroni corrected) and the sending matchup IBS (*t* (31) = 11.29, *p* < . 001, Cohen’s *d* = 1.91, Bonferroni corrected). Collectively, the results revealed that the IBS was only highly related to the recall quality of sharing emotional stories matchup (The Speaker Speaking-The Listener Listening).

### Neural-Behavioral correlation

A subsequent correlation analysis examined the association between the recall quality and the PLV in the theta band. 12 electrode pairs were found significantly higher PLV in the happy group than in the sad group. PLV in the theta band was aggregated to calculate the average PLV value of these 12 electrode pairs. Results revealed that the recall quality showed a significant, positive association with the PLV in the theta band in both happy and sad groups (*r_happy_* (15) = 0.61, *p_happy_* = 0.01, *r_sad_* (17) = 0.53,*p_sad_* = 0.03, Fig. 4A).

**Figure 4.**
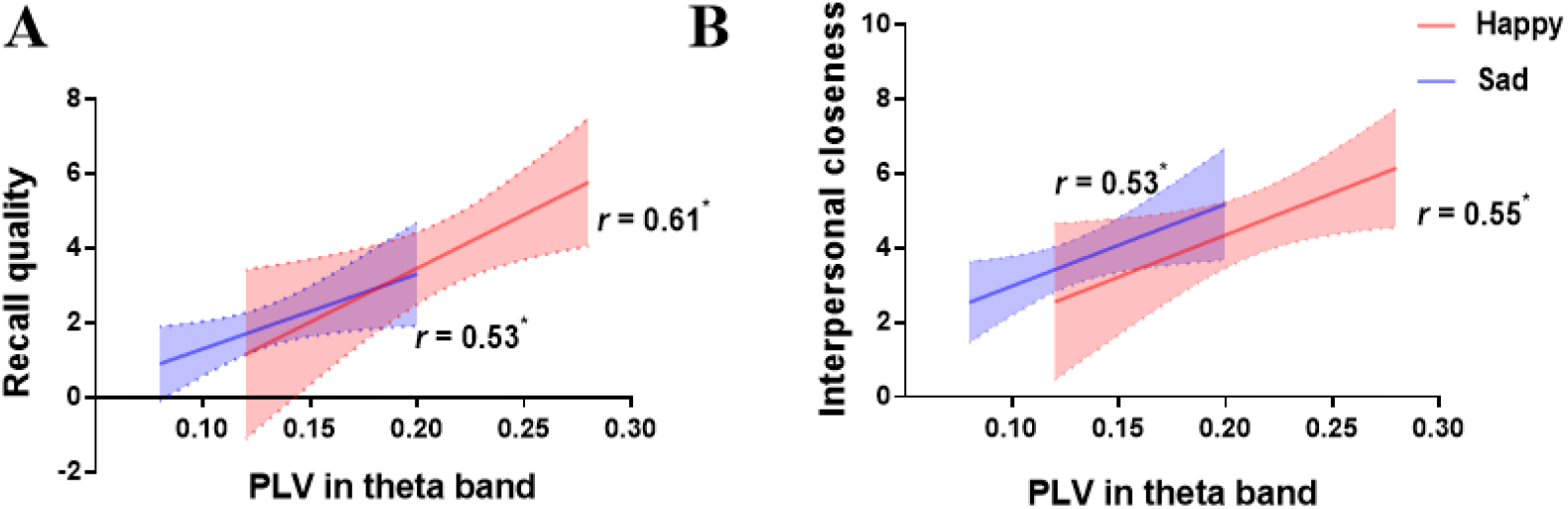
Correlation between behavioral results and IBS. (A) Correlation between the recall quality (delta value) and the PLV in the theta band (delta value). In both the happy group (Happy-Neutral) and sad group (Sad-Neutral), the differential theta band PLV (Happy/Sad minus corresponding Neutral) positively correlated with the differential recall quality of dyads (Happy/Sad minus corresponding Neutral). (B) Correlation between the interpersonal closeness and the PLV in the theta band. In both the happy group and sad group, the differential theta band PLV (Happy/Sad minus corresponding Neutral) positively correlated with the differential the interpersonal closeness of dyads (Happy/Sad group minus corresponding Neutral). \**p* < 0.05.

As for the relationship between subjective ratings and IBS, Pearson correlation analyses were conducted of differential theta band PLV and the differential interpersonal closeness in two emotional groups. Results revealed that the PLV in the theta band showed a significant, positive association with the ratings of the interpersonal closeness in both happy and sad groups (*r_happy_* (15) = 0.64, *p_happy_* = 0.01, *r_sad_* (17) = 0.58, *p_sad_* = 0.01, see Fig. 4B).

## Moderated mediation model

Previous results revealed: (a) a positive correlation between the recall quality and the PLV in the theta band in both happy and sad groups (Fig. 4A); (b) a positive correlation between the PLV in the theta band and the interpersonal closeness scores in both happy and sad groups (Fig. 4B). Therefore, it is plausible that IBS might mediate the association between the quality of sharing stories and interpersonal closeness. Further, emotion (Happy vs. Sad) might play a moderating role in this mediation model.

A moderated mediation model was estimated, revealing that recall quality positively predicts IBS (*β* = 0.26, *p* = 0.04), IBS positively predicts interpersonal closeness (*β* = 0.42, *p* < 0.01), and recall quality positively predicts interpersonal closeness (*β* = 0.43, *p* < 0.01). These results indicated a significant mediating effect of IBS in the relation between recall quality and interpersonal closeness. The interaction of recall quality for happy stories had a significant effect on the IBS (*β* = 0.16, *p* = 0.04), indicating that the relationship between recall quality and IBS was moderated by happy emotion (Fig. 5A). Namely, these results suggest that sharing happy stories can make speaker-listener dyads closer by increasing IBS. However, there was no significant moderated effect for sharing sad stories. Further, we conducted a simple slope test based on the moderated values at the mean and at ±1 standard deviation. The label (High/Low) of recall quality (delta value) was based on dichotomization via mean split. We found that the positive effect of recall quality on PLV in theta band is more pronounced in the happy emotion (see Fig. 5B).

**Figure 5.**
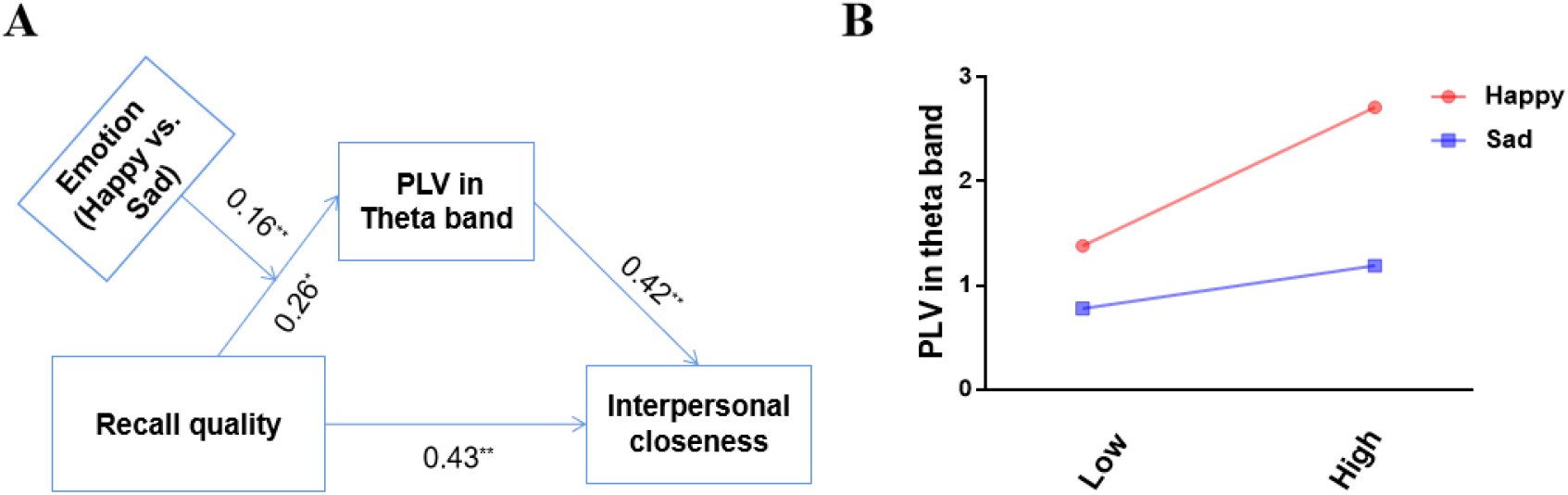
The moderated mediation model. (A) The mediation effect of theta-band interpersonal brain synchronization (delta value) on sharing stories (delta value) and interpersonal closeness (delta value) was moderated by happy emotion. (B) Simple slope test of the moderated effect of happy and sad group. The estimates presented here were standardized values. Low is the low recall quality (delta value), High is the high recall quality (delta value). \**p* < 0.05, \*\**p* < 0.01.

Taken together, the results demonstrated the mediating effect of IBS and the moderating effect of happy emotion in the relationship between the recall quality and the interpersonal closeness of speaker-listener dyads. Moreover, the moderated mediation model showed that the mediating effect of the IBS was influenced by the moderator, happy emotion.

### Coupling directionality

The G-causality analysis was used to measure the directional information flow (i.e., speaker → listener, listener → speaker). The paired-sample *t*-test showed that there was a significant difference between the two directions of G-causalities in The Speaker Speaking-The Listener Listening matchup (*t* (31) = 2.91, *p* < 0.01, Cohen’s *d* = 0.83; Fig. 6). To sum up, the result indicated that the speaker and the listener played an unequal role in the significant IBS during shared stories, that is, the information could only be transmitted from the speaker to the listener.

**Figure 6.**
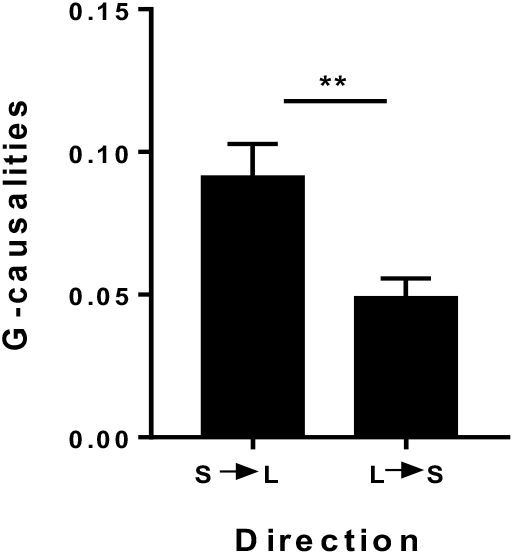
Directional coupling of the average PLV vaule of 12 significant electrodes pairs in theta band. G-causality direction from speaker to listener (S→L) and the reverse direction (L→S). There is a significant difference between the two G-causalities (S→L and L→S) at The Speaker Speaking-The Listener Listening matchup. S is speaker, L is listener; Error bars =±1 SEM; *\*\*p* < 0.01.

### Prediction of interpersonal closeness

Evaluating whether brain synchronization as a neural indicator in the present study can be used to predict the interpersonal closeness between speaker-listener dyads, we used SVC and SVR methods from machine learning. As expected, the AUC was high and statistically significant for distinguishing happy and sad emotions (AUC = 0.91, *p* = 1.4×10^-5^; Fig. 7A and B). Furthermore, results from the SVR analysis showed that the correlation coefficient between the actual and predicted interpersonal closeness of the testing dataset was approximately 0.67 (*p* = 1.13×10^-3^). Taken together, IBS can discriminate and predict the interpersonal closeness between speaker-listener dyads.

**Figure 7.**
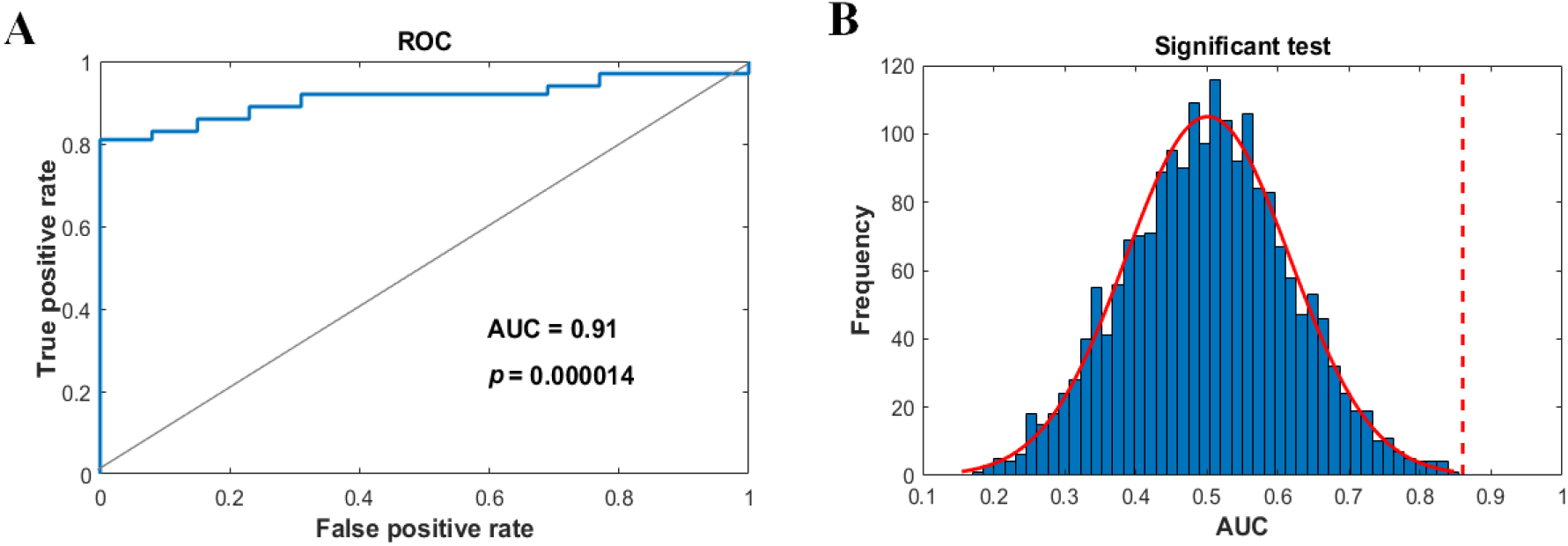
IBS discriminate and predict interpersonal closeness. (A) Receiver operating characteristic curve for classification distinguishing high rating of the interpersonal closeness from low interpersonal closeness. (B) The area under the receiver operating characteristic curve (AUC) is calculated as a metric of classification interpersonal closeness. The significance level (threshold at *p* < 0.05) is calculated by comparing the AUC from the correct labels (dotted line) with 2,000 randomization samples with shuffled labels (blue bars).

## Discussion

In the present study, we explored *(i)* the association between sharing stories and interpersonal closeness and (*ii*), its underlying neural correlates (*iii*) and, the different roles of happy and sad emotions (i.e., Happy vs. Sad) in sharing stories and building interpersonal closeness. As expected, the present study revealed that the behavioral index (the quality of recall after hearing an emotional story) was positively associated with interpersonal closeness in both happy and sad groups. Compared with the sad group, the happy group showed the better recall quality and reported higher interpersonal closeness. Moreover, higher task-related IBS was found for the happy group. Such IBS mediated the relationship between recall quality and interpersonal closeness. Within the happy group, sharing stories moderated the IBS and thus promoted interpersonal closeness. Finally, the IBS can be used to predict interpersonal closeness. The aforementioned results are discussed in detail as follows.

Our results showed a positive association between sharing emotional stories and interpersonal closeness at the behavioral level. In line with the previous observations regarding the benefits of sharing information on the interpersonal relationship, sharing stories can increase mutual understanding and promote communication between individuals (Dubois et al., 2016; McCarthy & Duck, 1976). Further, in the light of EASI, humans have the tendency to synchronize with each other’s behavior (Dimberg & Thunberg, 1998) and physiological states (Konvalinka et al., 2011) during emotional expression. In line with EASI, our results suggest that sharing happy and sad stories can facilitate interpersonal closeness. Positive emotional information is more likely to be transferred and received (Piotroski, Wong, & Zhang, 2015) by representing a positive image (Birnbaum, Iluz, & Reis, 2020). On the behavioral level, this result further contributed to the reconciliation of the controversy on the effect of sharing different emotional stories on interpersonal closeness. Specifically, our findings show that, compared with the sad group, the happy group showed the better recall quality and led to higher interpersonal closeness.

Examining the cognitive and neural processes involved in social interaction behaviors hinges on investigating brain-to-brain synchronization during social interaction. “Two-person neuroscience” in sharing stories has higher ecological validity than single-brain recoding because “two-person neuroscience” is closer to the interaction in real life (García & Ibáñez, 2014; Joy, Adam, Zhang, Swethasri, & Yumie, 2018; Redcay & Schilbach, 2019). Moreover, brain-to-brain studies have been widely accepted to unveil the interpersonal neural correlates in the context of social interactions (Chen et al., 2020; Lu, Xue, Nozawa, & Hao, 2018). Therefore, the present study used the brain-to-brain recording to evaluate the dynamic neural interaction between the speaker and the listener, revealing the brain-to-brain interaction pattern in the process of sharing stories. Referred to previous sharing emotional stories brain-to-brain studies and audio brain-to-brain studies (e.g., Hou et.al, 2020; Smirnov et.al, 2019), the present study was not real-time interction.

We found significant IBS between speaker and listener in the frontal and temporal cortices during interaction. Previous brain-to-brain studies have found strong interpersonal neural synchronization in the frontal cortex and the temporal cortex by using the interactive paradigm involving verbal communication (Ahn et al., 2018; Bohan et al., 2018). First, prior studies have uncovered that the frontal cortex critically contributes to recognize emotion and encode information (Abrams et al., 2010; De Borst, Valente, Jääskeläinen, & Tikka, 2016). Moreover, human neuroimaging studies have implicated the temporal area in the high-level metallization during communications (Samson, Apperly, Chiavarino, & Humphreys, 2004). Therefore, our finding is consistent with previous findings, demonstrating that the listener needs to accurately understand the emotion of the stories and establish a frame of information related to the stories which are related to frontal cortex and temporal cortex.

Our results indicated the IBS played a mediating role between sharing stories and interpersonal closeness. Based on the neuroimaging studies, sharing stories may be inherently reflected in the neural level (Berger & Jonah, 2014; Tamir & Mitchell, 2012), and comprehension of narrations was driven by the neural similarity between the speaker and the listener (Nguyen et al., 2019; Silbert, Honey, Simony, Poeppel, & Hasson, 2014). Similar understanding of stories during interpersonal interaction leads to enhanced IBS which represented the higher neural similarity of speaker-listener dyads (Chen et al., 2020; Hasson, Ghazanfar, Galantucci, Garrod, & Keysers, 2012; Jiang et al., 2015; Nozawa, Sasaki, Sakaki, Yokoyama, & Kawashima, 2016). Consistent with previous studies (Kulahci & Quinn, 2019; Tamir & Mitchell, 2015), our findings suggested that high IBS levels represented the high-level understanding of sharing stories and it was essential to increase interpersonal closeness between individuals.

Our findings also pointed to the moderation effect of happy emotion between sharing stories and the IBS. A recent study has shown that the neural synchronization between the speaker and the listener was associated with the emotional features of stories and the neural synchronization created a tacit understanding between the speaker and the listener, facilitating communication and improving interpersonal relationships (Smirnov et al., 2019). It is worth noting that only the happy emotion (relative to neutral) played a moderated role in enhancing IBS. Although the theory of EASI proposes that emotional expression will increase mutual understanding between individuals, compared with positive emotional expression, the effect of negative emotional expression is worse (Dubois et al., 2016). As demonstrated by previous studies, bragging about their positive information can make people feel good and improve interpersonal relationships by increasing activation in the nucleus accumbent and ventral tegmental area, which form the mesolimbic dopamine system (Tamir & Mitchell, 2012). Individuals prefer to share positive self-related details (i.e., happy stories about the videos you watched) in the presence of the strangers and they also wish to transfer an ideal image (Baek et al., 2017; Tamir & Mitchell, 2012). On the neural level, our result further contributed to the role of different emotions (i.e., Happy vs. Sad) in the mediation effect of sharing different stories on interpersonal closeness. To sum up, our results supported that sharing happy stories, instead of sharing sad stories, were useful for enhancing speaker-listener interaction.

Our GCA results further showed that there was a significant directionality of the enhanced IBS between the speaker and the listener, implying that the speaker and the listener displayed a differential and asymmetrical role during sharing emotional stories. Our findings were consistent with previous studies in unilateral communication or unilateral sharing, the speaker owns more information than the listener (Tworzydło, 2016). Based on the verbal cues of the speaker, the listener would frame the information, fill in the content, and adjust the content during the dynamic interactive process (Chen et al., 2017; Nguyen et al., 2019; Zadbood et al., 2017). In line with previous findings, listeners would perform as followers during sharing emotional stories and this performance is influenced by the speaker (Bohan et al., 2018; Jiang et al., 2015; Stephens et al., 2010). Thus, the importance of the speaker and the listener were different in the sharing process, and the directionality of IBS in our study highlighted the point that sharing emotional stories was dominated by the speaker.

Our results of the discriminant analysis indicated that the IBS of speaker-listener dyads was able to distinguish the rating of the interpersonal closeness through SVC, and further revealed the prediction of IBS for interpersonal closeness through SVR. These findings were in line with recent studies revealing that synchronized brain activity served as a reliable neural-classification feature (Cohen et al., 2018; Hou et al., 2020; Pan et al., 2020). Moreover, a growing number of studies have used the combination of machine learning and the IBS measurement in social neuroscience, so we considered more features, such as the Time-frequency neural features from single-trial event-related spectral perturbation (ERSP) patterns (Chung, Yun, & Jeong, 2015; Scott Makeig, Debener, Onton, & Delorme, 2004).

The present study has several limitations. First of all, spatial resolution is restricted in EEG which is distributed on by the skull and scalp (Hedrich, Pellegrino, Kobayashi, Lina, & Grova, 2017), limiting measurements to specific areas during sharing emotional stories of speaker-listener dyads. Our findings indicated that frontal and temporal cortices were important in sharing emotional stories. Although the ventromedial prefrontal cortex (VMPFC) and anterior cingulate cortex (ACC) play a crucial role in sharing emotional information activities (Killgore et al., 2013), EEG is unable to measure these two areas. Second, the present study focused on specific stimuli (one happy story and one story). Thus, future work is encouraged to use a larger range of stimuli to generalize these results to happy and sad stories more broadly beyond the specific stimuli used here. Finally, the exploratory SVC/SVR predictive analysis is constrained by a relatively sample size (although our sample size is similar to those reported in previous classification analyses based on brain-to-brain coupling data; e.g., Dai et al., 2018; Jiang et.al, 2012; Pan et.al, 2020). Future replications are encouraged to consolidate the current findings by increasing both the sample size and number of testing blocks.

To conclude, the present study showed that sharing both happy and sad stories could increase interpersonal closeness between individuals. Moreover, findings at the neural level suggested that only sharing happy stories moderated the IBS and thus promote interpersonal closeness. These insights contribute to a deeper understanding of the neural correlates of sharing different emotional stories with interpersonal closeness. Future research may explore the neural mechanism of sharing stories by using IBS as an effective neural indicator.

## Acknowledgments

This work was supported by the Shanghai Key Base of Humanities and Social Sciences (Psychology-2018), the National Natural Science Foundation of China (32071082 and 71942001), Key Specialist Projects of Shanghai Municipal Commission of Health and Family Planning (ZK2015B01), and the Programs Foundation of Shanghai Municipal Commission of Health and Family Planning (201540114).

