## Supplements for "Sharing happy stories increases interpersonal closeness mediated by enhancing interpersonal brain synchronization"

### Supplementary materials

**Table S1.** Example of one rater. An example of a rater's behavioral rating sheet. The rater has specified 1, 0.5, or 0 points respectively for fully describing, only mentioning, or not mentioning each of 20 important scenes. The rater has included an overall comprehension level out of 10 and the total score for each subject was out of 30. Overall high correlation between the ratings of different raters validated this approach.

| scene name | S1 | S2 | S3 | S4 | S5 | S6 | S7 | S8 | S9 | S10 | S11 | S12 | S13 | S14 | S15 |
|---|---|---|---|---|---|---|---|---|---|---|---|---|---|---|---|
| This is a happy story | 1 | 1 | 1 | 1 | 1 | 1 | 1 | 1 | 1 | 1 | 1 | 1 | 1 | 1 | 1 |
| Case will be retrial | 1 | 0 | 0 | 1 | -1 | 0 | 0 | 0 | 0 | 1 | 0 | 0 | 0 | 0 | 0 |
| Lord Bao was teased about Tan | 0.5 | 0 | 0 | 0 | 1 | 0.5 | 1 | 0 | 0.5 | 0.5 | 0 | 0.5 | 0 | 0 | 0 |
| Three bosses appeared | 0.5 | 0.5 | 1 | 1 | 1 | 0 | 1 | 1 | 1 | 1 | 1 | 1 | 1 | 0.5 | 1 |
| Explained the rules | 0 | 0 | 0.5 | 0.5 | 1 | 1 | 0.5 | 0.5 | 0 | 1 | 0 | 0 | 0.5 | 0 | 0.5 |
| Preparation before the debate | 0 | 1 | 0 | 1 | 1 | 1 | 1 | 1 | 1 | 1 | 0 | 0.5 | 0.5 | 0.5 | 0.5 |
| Three bosses in cahoots | 0 | 0.5 | 0 | 0 | 0 | 0 | 0 | 0 | 0 | 0 | 0 | 0 | 0 | 0 | 0 |
| Three bosses gave the signal | 0 | 0 | 0 | 0 | 0 | 0 | 0 | 0 | 0 | 0 | 0 | 0 | 0 | 0 | 0 |
| Lord Bao made some abusive remarks | 1 | 1 | 1 | 1 | 1 | 1 | 1 | 1 | 1 | 1 | 1 | 0 | 1 | 0 | 1 |
| Lord Bao took a swipe at bosses | 0 | 0 | 0 | 0 | 0 | 0 | 0.5 | 0 | 0 | 0 | 0 | 0 | 0 | 0 | 0 |
| The reason Lord Bao made abusive remarks | 0.5 | 0 | 0.5 | 1 | 0.5 | 0 | 1 | 0 | 0 | 0.5 | 0 | 0 | 0 | 0 | 0 |
| A sidekick fell down | 1 | 1 | 0 | 0 | 0.5 | 1 | 1 | 1 | 1 | 1 | 0 | 0 | 0 | 0 | 0 |
| Lord Bao asked the defendant to stand back | 0 | 1 | 1 | 0.5 | 0.5 | 0.5 | 0 | 0 | 0.5 | 1 | 0.5 | 0.5 | 0.5 | 0.5 | 0 |
| Lord Bao fought with the defendant | 1 | 1 | 0 | 1 | 1 | 0 | 1 | 0.5 | 0.5 | 1 | 1 | 0.5 | 1 | 0 | 0 |
| Lord Bao is arguing with his bosses | 0 | 0 | 0 | 0 | 0 | 0 | 0 | 0 | 0 | 0 | 0 | 0 | 0 | 0 | 0 |
| Here comes the prisoner | 1 | 0.5 | 0 | 1 | 1 | 0 | 1 | 0.5 | 0 | 1 | 0.5 | 1 | 0.5 | 0 | 1 |
| Lord Bao argued with the prisoner | 1 | 0 | 0 | 1 | 1 | 0.5 | 1 | 0 | 1 | 1 | 1 | 0 | 1 | 0 | 1 |
| Lord Bao criticized the prisoners | 0 | 0 | 0 | 0 | 0 | 0 | 0 | 0 | 0 | 0 | 0 | 0 | 0 | 0 | 0 |
| People dress up as dogs | 0 | 1 | 1 | 1 | 1 | 0 | 1 | 0 | 0.5 | 0 | 0 | 1 | 1 | 0 | 1 |
| Lord Bao won the retrial | 0 | 0.5 | 0 | 1 | 0.5 | 0.5 | 0 | 0 | 0 | 0.5 | 0 | 0.5 | 0 | 0 | 0 |
| sum of scene scores out of 20 | 8.5 | 9 | 6 | 12 | 11 | 7 | 12 | 6.5 | 8 | 12.5 | 6 | 6.5 | 8 | 2.5 | 7 |
| overall memory out of 10 | 6 | 6 | 5 | 7 | 4 | 4 | 6 | 5 | 6 | 7 | 6 | 5 | 6 | 2 | 5 |
| score (out of 30) | 14.5 | 15 | 11 | 19 | 15 | 11 | 18 | 11.5 | 14 | 19.5 | 12 | 11.5 | 14 | 4.5 | 12 |

**Figure S1.** Increased inter-brain PLVs. The electrodes that showed significantly increased inter-brain PLVs with Happy (vs. Sad) brain-to-brain coupling in other bands (all Bonferroni corrected).

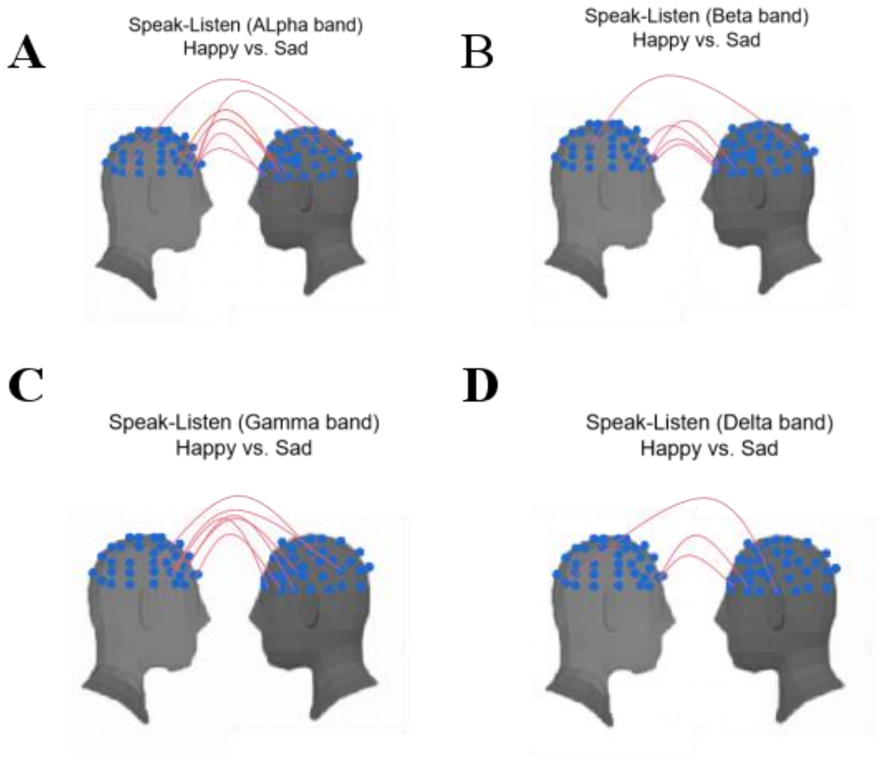

**Table S2.** Specific links of speaker-listener dyad.

| Speaker | Listener | *t* value | *P* value (Bonferroni corrected) |
|---|---|---|---|
| FC4 | F5 | 7.90 | 4.15E-10 |
| FC2 | FC5 | 3.98 | 1.28E-05 |
| FPZ | FC5 | 7.62 | 8.24E-10 |
| FC6 | FC5 | 3.56 | 3.93E-05 |
| F2 | F5 | 5.32 | 3.57E-07 |
| FC2 | F5 | 4.16 | 8.63E-06 |
| FPZ | F5 | 5.73 | 1.20E-07 |
| C2 | FPZ | 3.94 | 1.42E-05 |
| FC2 | F7 | 4.38 | 4.76E-06 |
| FP2 | FT7 | 9.67 | 6.34E-12 |
| FP2 | F7 | 3.53 | 4.24E-05 |
| O2 | AF3 | 4.03 | 1.21E-05 |

**Figure S2.** The average PLV values of 12 significant electrodes pairs in theta band in different stages. PLV in theta band (delta value) is shown for happy (Happy-Neutral) and sad (Sad-Neutral) emotions and compared sharing stage (The Speaker Speaking-The Listener Listening) was compared to the receiving stage (The Speaker Watching-The Listener Listening) and sending stage (The Speaker Speaking-The Listener Recalling). Error bars = ± 1 SEM; $^{***}p < 0.001$, after Bonferroni corrected.

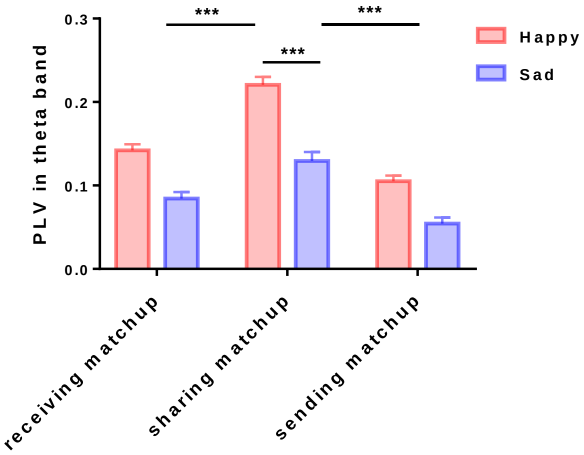
